# The preference for surprise in reinforcement learning underlies the differences in developmental changes in risk preference between autistic and neurotypical youth

**DOI:** 10.1101/2024.08.23.609310

**Authors:** Motofumi Sumiya, Kentaro Katahira, Hironori Akechi, Atsushi Senju

**Affiliations:** Research Center for Child Mental Development, Hamamatsu University School of Medicine, Hamamatsu, Japan; United Graduate School of Child Development, Osaka University, Kanazawa University, Hamamatsu University School of Medicine, Chiba University and University of Fukui, Osaka/Kanazawa/Hamamatsu/Chiba/Fukui, Japan; Human Informatics and Interaction Research Institute, National Institute of Advanced Industrial Science and Technology (AIST), Tsukuba, Japan; Graduate School of Comprehensive Human Sciences, University of Tsukuba, Tsukuba, Japan; Graduate School of Education, Kyoto University, Kyoto, Japan

**Author notes:** Corresponding author, Motofumi Sumiya, PhD, (M.S.).

**Keywords:** risk preference, developmental changes, autism, reinforcement learning, surprise

## Abstract

Risk preference changes nonlinearly across development. Although extensive developmental research on the neurotypical population has shown that risk preference is highest during adolescence, developmental changes in risk preference in autistic people, who tend to prefer predictable behaviors, have not been investigated. Here, we aimed to investigate these changes and underlying computational mechanisms. Using a game-like risk-sensitive reinforcement learning task, we found a significant difference in nonlinear developmental changes in risk preference between the autistic and neurotypical groups (N = 75; age range, 6–30 years). The computational modeling approach with reinforcement learning models revealed that individual preferences for surprise modulated such preferences. These findings indicate that for neurotypical people, adolescence is a developmental period involving risk preference, possibly due to lower surprise aversion. Conversely, for autistic people, who show opposite developmental trajectories of risk preference, adolescence could be a developmental period involving risk avoidance because of low surprise preference.

## Introduction

Risk preference, the propensity to take or avoid risk, is a fundamental driver of decision-making, which reportedly undergoes complex developmental changes across one’s lifespan. Psychometric questionnaire surveys and laboratory task studies provide empirical evidence supporting a developmental peak in risk-taking in mid-adolescence, that is, nonlinear developmental changes in risk-taking that increase and decrease after adolescence (1–5). Conversely, adolescents are also known to make more prudent decisions than adults, depending on the task and context (6–8). Although extensive research on neurotypical (NTP) individuals has revealed the developmental trajectory of changes in risk preference, our understanding of how these processes unfold in autistic (AUT) individuals remains limited.

Clinically, AUT is defined as a neurodevelopmental condition that is primarily characterized by difficulties in social communication and interaction, as well as restricted, repetitive patterns of behavior, interests, or activities (9). In recent years, multiple studies with large numbers of research participants have shown that everyday risky behaviors, such as binge drinking or using illicit drugs, which are considered problematic in NTP individuals, are less common in AUT individuals (10, 11). Such risk preferences in AUT individuals have also been investigated in experimental studies. Recently, van der Plas et al. (12) conducted a narrative review of 104 decision-making studies on AUT people and found that their performance in reward-learning paradigms (e.g., learning which deck of cards provides the best reward) was similar to that of NTP individuals, but their performance in value-based paradigms (e.g., making a decision based on a choice between two outcomes that differ in subjective value) was different from that of NTP individuals. For example, in several studies that adopted value-based paradigms, financial risk-taking tasks focused on choices between alternatives of equivalent objective value but different risk for rewards (i.e., a choice between a sure option to win $1 versus a risky option to have a 50% chance to win $2) have shown that AUT adults are more risk-averse than NTP adults (13–15). Although these studies suggest that differential subjective value processing, not reward learning, in risk-taking decisions underlies the risk aversion in AUT people, the core underlying mechanism that affects subjective decision-making is not known.

The computational modeling approach has helped us investigate the possible computational mechanisms underlying risk-taking decisions primarily in the NTP population. Multiple reinforcement learning models, such as the utility model, which incorporates nonlinear subjective utilities for different amounts of reward, or risk-sensitive model in which positive and negative prediction errors (differences between a decision outcome and its predicted outcome) have asymmetric effects on learning, have been proposed to explain risk-taking decision-making (16, 17). Recently, we proposed that the preference for surprise in a decision outcome is a critical factor that can modulate risk preference; surprise occurs because of prediction errors, regardless of whether the error is positive or negative (18). Based on the cognitive evolutionary model of surprise (19) and experimental studies (20–22), we proposed a reinforcement learning model in our previous study——the surprise model——by introducing a parameter that modulates the value of the outcome based on surprise in each decision (18). Using two datasets of risk-taking tasks with monetary outcomes (16, 23) and behavioral simulations, we showed that the surprise model had a better fit, indicating that surprise in each decision leads individuals to risk-averse behavior. Therefore, we hypothesized that the tendency toward risk aversion in AUT people is explained by their aversion to surprise. This is because AUT individuals tend to have a strong preference for routine, and repetitive behaviors are hypothesized to stem from the different ways in which prediction errors are processed, which could lead to a preference for predictable behaviors or situations (23, 24). Accordingly, it is possible that the preference for surprise, as a core mechanism underlying the developmental change in risk preference, leads to a difference in risk preference between the two neurodiverse populations.

In this study, we aimed to investigate the nonlinear developmental changes in risk preference in AUT and NTP individuals, as well as to elucidate the underlying computational mechanism from the perspective of surprise preference, by adopting a multi-method approach that integrated online experimental paradigms, cross-sectional data, and computational modeling. We hypothesized that the risk preferences (i.e., choices between alternatives with the same objective value but different risks) of AUT people would differ from those of NTP people, which change nonlinearly across development. Furthermore, we hypothesized that this difference in risk preference between the two neurodiverse populations would be underpinned by a preference for surprise.

To test the hypotheses, we recruited 125 participants aged 6–30 years, and after multiple data exclusion steps, we analyzed the data of 75 participants (31 in the AUT group and 44 in the NTP group). For this study, we created a treasure task based on a risk-sensitive reinforcement learning task introduced by Rosenbaum et al. (17) (Fig. 1), without the use of monetary rewards to delineate decisions in many aspects of life. This is because decisions for monetary rewards may follow a developmental trajectory different from that of other types of risk-taking (17, 25, 26), which could confound our results. Focusing on choices between alternatives with the same objective value but different risks, we calculated the risk preference and stay probability of a risky choice after a rewarding or non-rewarding outcome. Analyses with t-tests and multiple regression analyses were conducted. Using the choice-related data of each participant, we fit four reinforcement learning models, three models that were previously used in a similar experimental paradigm (16, 17) and the surprise model (18), and compared the fit of each model to the data. Furthermore, we validated the results of model fitting with multiple methods, model recovery, parameter recovery, and posterior predictive check (27, 28).

**Fig. 1.**
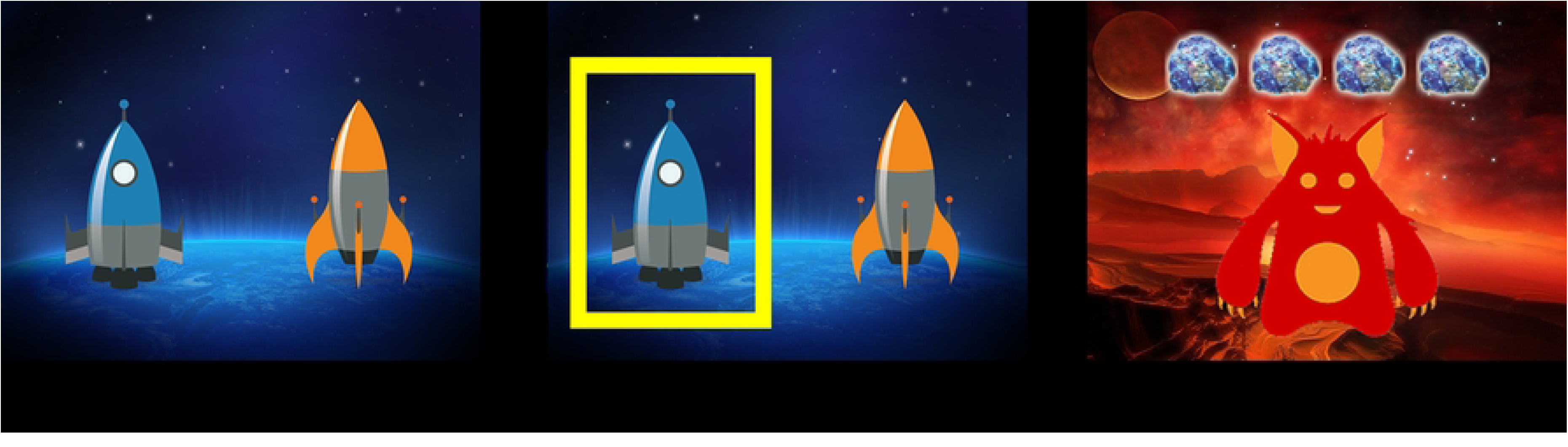
Task sequence.

## Results

### Risk preference

We did not find a significant group difference in risk preference (*t*=0.219, *p*=0.827). On the other hand, the multiple regression analysis revealed a significant interaction of the group and quadratic term of age (*β*=0.098, *p*=0.023) (Fig. 2a). The results of the other regressors are summarized in Fig. 3 and S2 Table. We calculated the multicollinearity of the regressors and confirmed weak correlations between them (variance inflation factor (VIF) <5) (S1a Fig.).

**Fig. 2.**
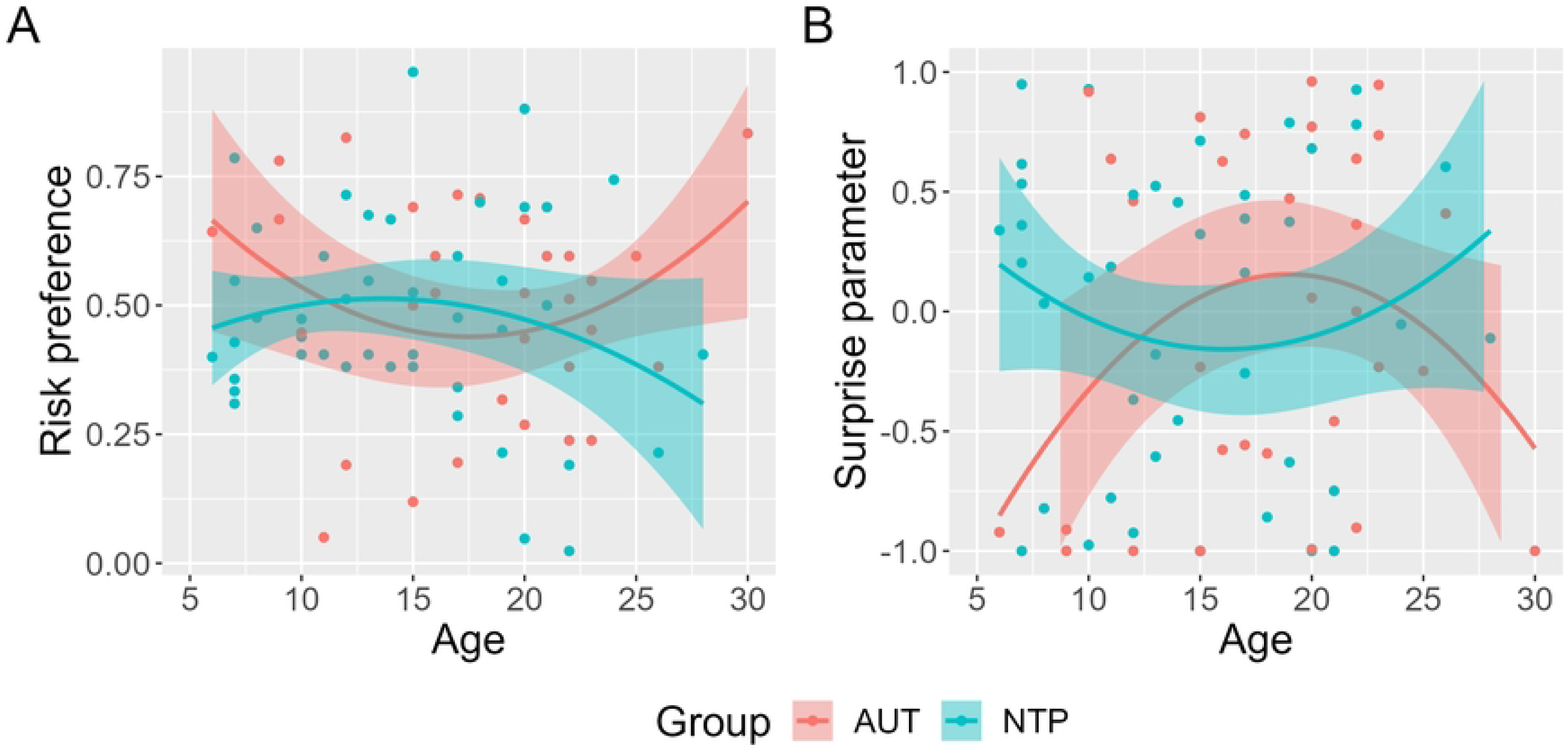
Relationship of age with (a) the risk preference and (b) estimated surprise parameter. The regression lines are from a linear regression model including linear and quadratic age terms. Data points represent individual participants. The lines and 95% confidence intervals were estimated under heterogeneous variances to reduce the influence of outlier data.

**Fig. 3.**
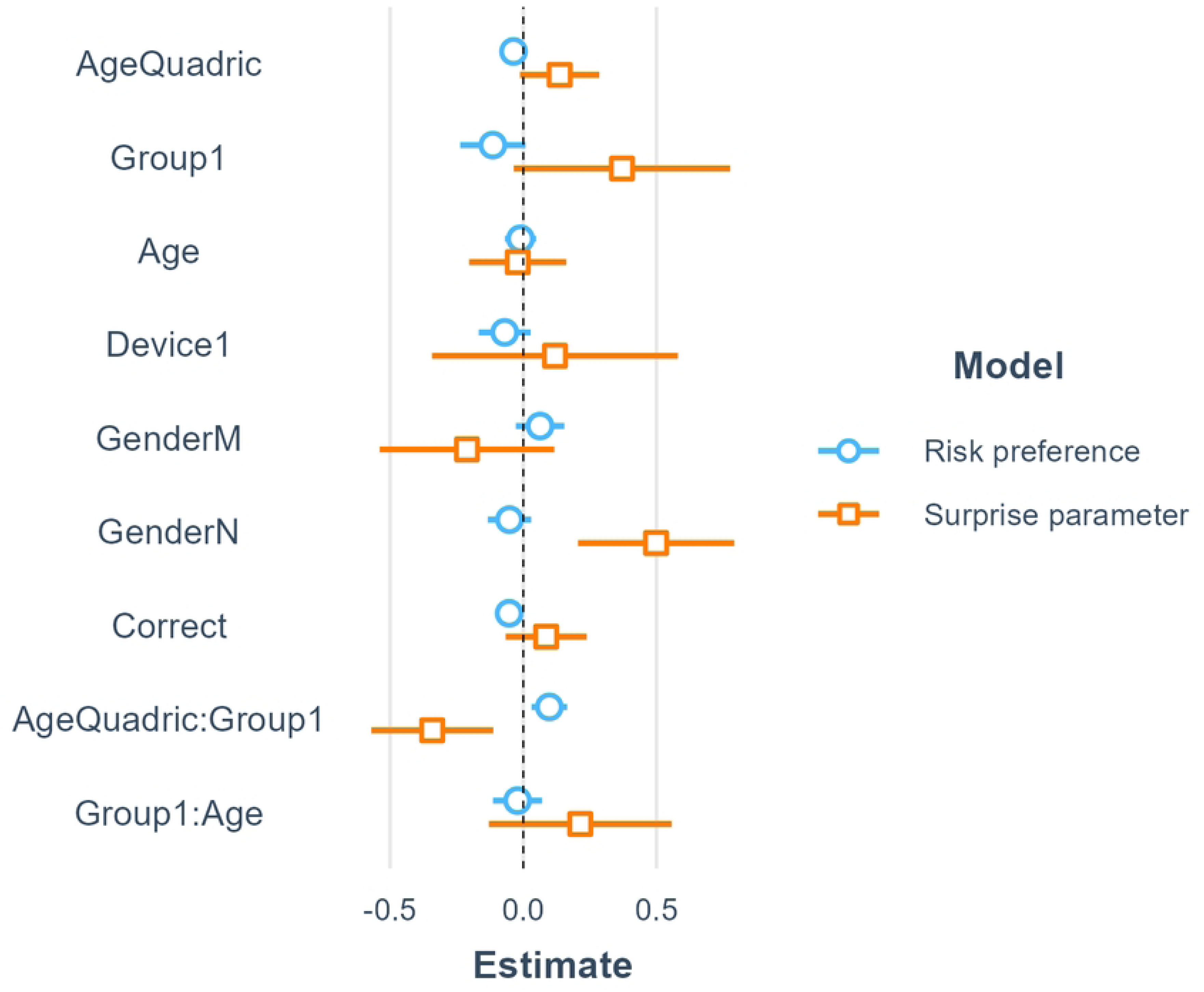
Model coefficients for risk preference (blue) and the surprise parameter (orange). Parameters within models were coded as group (AUT = 1, NTP = −1), device (PC = 1, Tablet = −1), and gender (M = male, F = female, N = not reported).

With consecutive multiple regression analyses within each group, we confirmed the tendency of an inverted U-curve developmental change in the NTP group (*β*=-0.039, *p*=0.059) and a U-curve developmental change in the AUT group (*β*=0.055, *p*=0.060) (S2 and S3 Fig., S3 Table).

### Estimated parameters

We used the surprise parameter of the surprise model, the winning model in the model comparisons described below, as the dependent variable in the multiple regression analysis. We found a significant interaction between the parameter and quadratic term of age (*β*=-0.342, *p*=0.017) (Fig. 2b). The results of the other regressors are summarized in Fig. 3 and S2 Table. Information on the multicollinearity of the regressors is summarized in S1b Fig.

With consecutive multiple regression analyses within each group, we confirmed the significance of a U-curve developmental change in the NTP group (*β*=0.142, *p*=0.046) and an inverted U-curve in the AUT group (*β*=-0.203, *p*=0.034) (S4 and S5 Fig., S4 Table).

In addition to the surprise parameter, we conducted multiple regression analysis with the other parameters in the surprise model: learning rate and inverse temperature. To address multicollinearity, we only modeled the interaction term of the group and age and of the group and quadratic term of age. The effect of the interaction between the group and quadratic term of age was significant on the inverse temperature (*β*=-1.871, *p*=0.025) but not the learning rate (*β*=--0.029, *p*=0.384) (S6, S7, and S8 Fig., S5 Table). These data are consistent with that in previous literature on the NTP population (29–31) and have added new findings about the AUT population (S1 Text).

### Stay probability

We performed a multiple regression analysis for each stay probability for the sure and risky choices. We found a significant interaction of the group and quadratic term of age for the stay probability of sure choices (*β*=-0.092, *p*=0.025) (Fig. 4a). For the stay probability of risky choices after a non-rewarding outcome, we removed the covariates——accuracy, gender, and device—from the model to address the excessive correlation among the explanatory variables. We found a significant interaction of the group and quadratic term of age (*β*=0.094, *p*=0.034) (Fig. 4b). Furthermore, we found no significant relationship but only a tendency toward one between the group and quadratic term of age for the stay probability of risky choices after a rewarding outcome (*β*=0.056, *p*=0.077) (Fig. 4c). The results for the other regressors are summarized in Fig. 5 and S6 Table. Information on the multicollinearity of the regressors is summarized in S9 Fig.

**Fig. 4.**
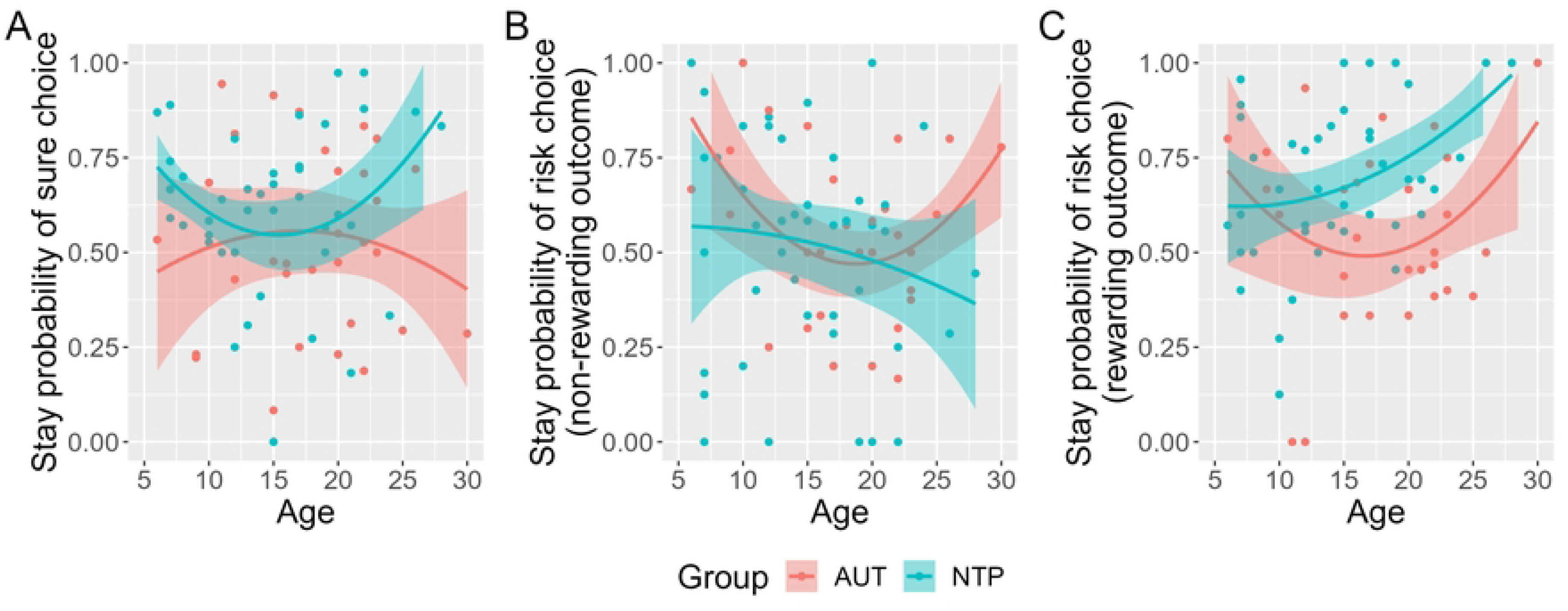
Relationship between age and the stay probability of a (A) sure choice, (B) risky choice after a non-rewarding outcome, and (C) risky choice after a rewarding outcome. The regression line is from a linear regression model including linear and quadratic age terms. Data points represent individual participants. The lines and 95% confidence intervals were estimated under heterogeneous variances to reduce the influence of outlier data.

**Fig. 5.**
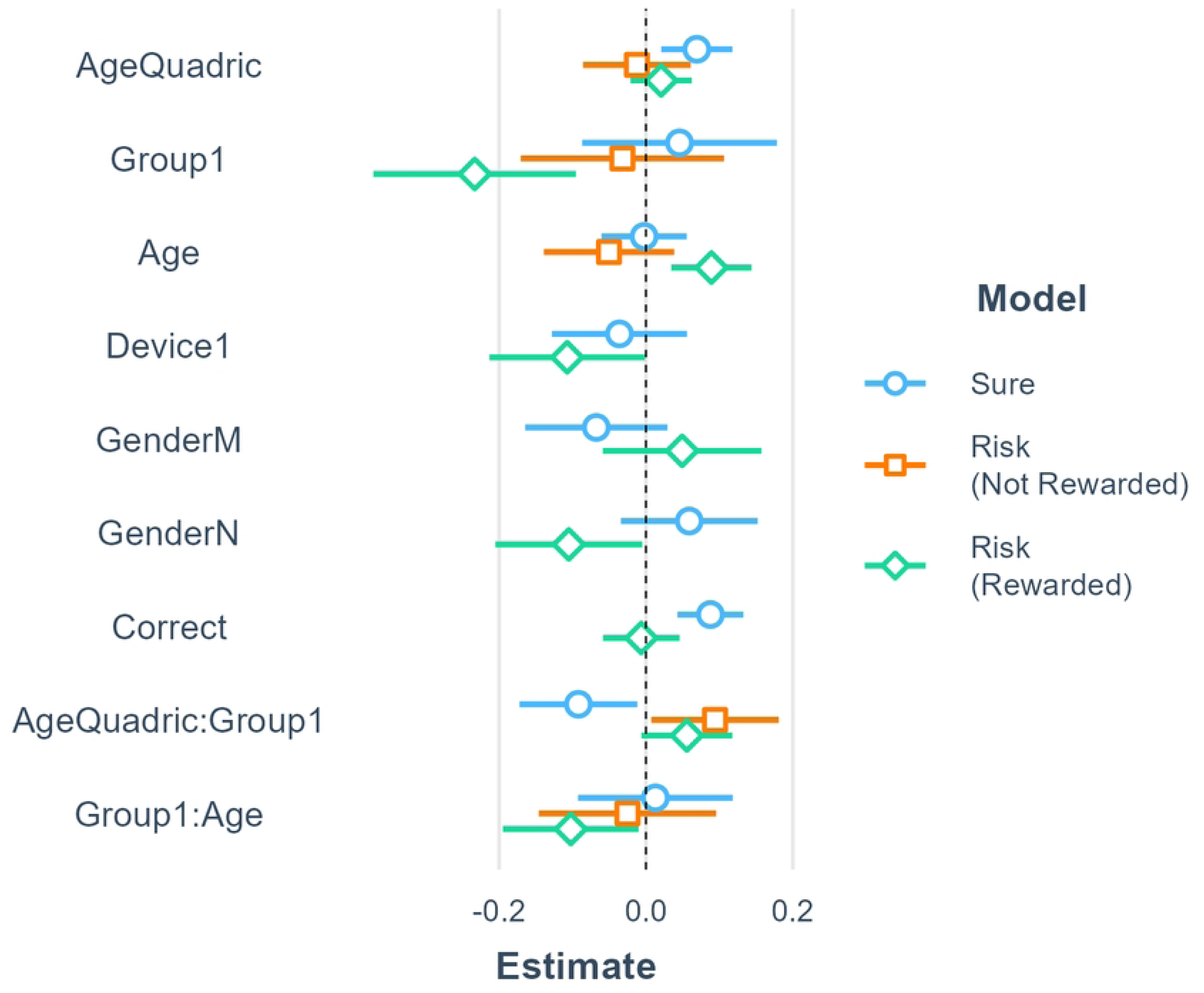
Model coefficient for stay probability for a risky choice after a non-rewarding outcome (blue) and rewarding outcome (orange). Parameters within models were coded as group (AUT = 1, NTP = −1), device (PC = 1, Tablet = −1), and gender (M = male, F = female, N = not reported).

We conducted consecutive multiple regression analyses for each stay probability within each group using the same regressors as those in the risk preference analysis in each group, but in the analysis of the stay probability for sure choices in the NTP group, a regressor (Device) was removed to deal with multicollinearity. We confirmed the significance of a U-curve developmental change in sure choices in the NTP group (*β*=0.071, *p*=0.004) but not in the AUT group (*β*=-0.011, *p*=0.558). Further, we confirmed the significance of a U-curve developmental change in risky choices after a non-rewarding outcome in the AUT group (*β*=0.080, *p*=0.002) but not in the NTP group (*β*=-0.016, *p*=0.673). Furthermore, we confirmed the significance of a U-curve developmental change in risky choices after a rewarding outcome in the AUT group (*β*=0.073, *p*=0.010) but not in the NTP group (*β*=0.020, *p*=0.316). Instead, the linear effect of age on risky choices after a rewarding outcome was significant in the NTP group (*β*=0.072, *p*=0.007) (S10 and S11 Fig., S7 Table).

### Model comparison

We compared the model evidence for each model (log marginal likelihood) and found that the surprise model had the highest value (Fig. 6a). We then performed a Bayesian model comparison to determine the best model to explain choice behavior and found that the surprise model had a significantly higher protected exceedance probability than the other models, indicating that it was more frequent in this population (Fig. 6b). We further checked the fitness of each model in each group and confirmed that the surprise model had the best fit among the four models (S12a-d Fig.). We also confirmed that the fit of the surprise model was the best in a relatively large proportion of participants (42% in the AUT group, 41% in the NTP group) compared to that of the other models (utility model: 35% in the AUT group, 20% in the NTP group; Q-learning [QL] model: 13% in the AUT group, 27% in the NTP group; risk-sensitive [RS]QL model: 9.7% in the AUT group, 11% in the NTP group) (S8 Table). The distribution of each estimated parameter in the surprise model is shown in S13 Fig. to illustrate the potential problem fitting (27, 32). These results are discussed in S2 Text.

**Fig. 6.**
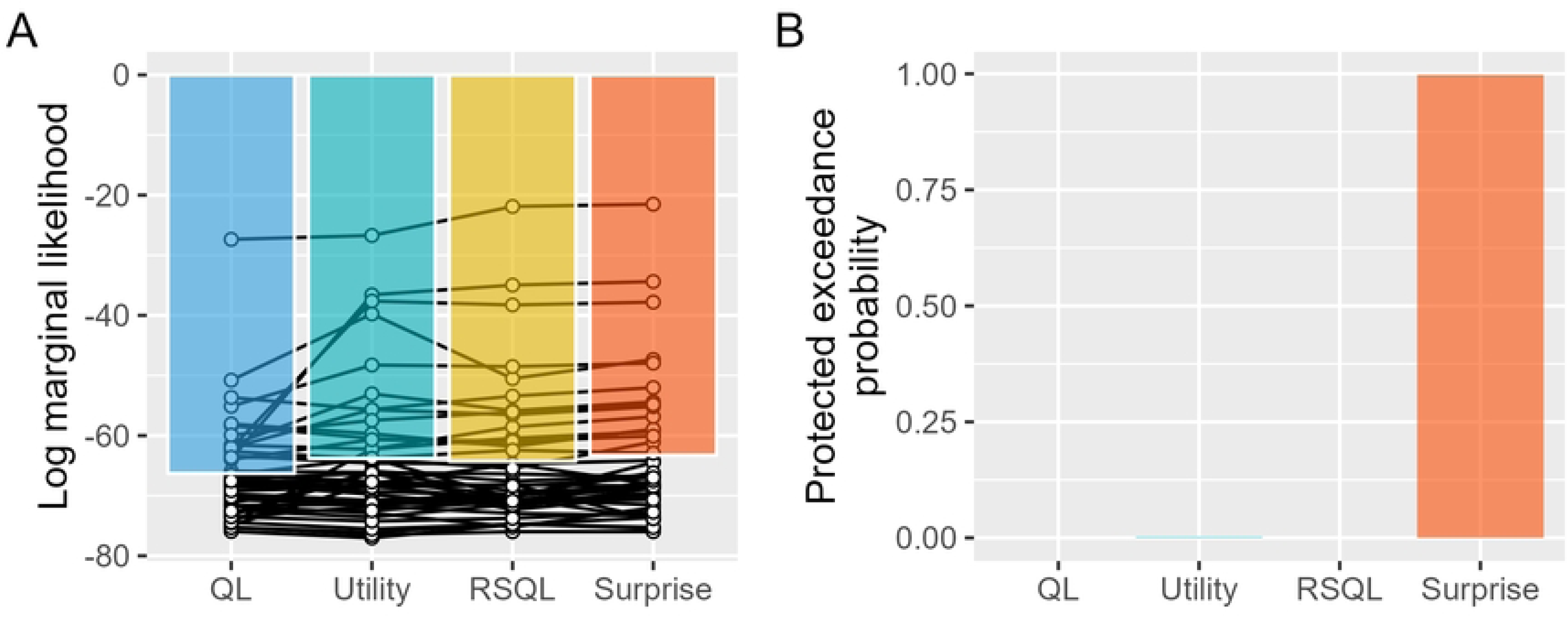
Model comparison. (a) Evidence of each model (log marginal likelihood). (b) Bayesian model comparison.

### Parameter recovery

For most of the parameters in all models, the recovered and true parameters were highly correlated (*r* >0.91) (QL: alpha, utility: alpha & utility, RSQL: alphaP & alphaN, surprise: alpha & surprise), confirming that these parameters were identifiable (S9 Table). These findings indicate that the model parameters can be adequately estimated for all models and that the estimated parameters are meaningful.

### Model recovery

S14 Fig. shows the confusion matrix resulting from the model recovery process. The numbers denote the frequencies at which each fitted model was selected for each simulated model. The simplest model, the QL model, exhibited a high recovery rate (89%). As complex models, the utility and surprise models showed similarly high recovery rates to those of the QL model (utility: 54%, surprise: 53%), which indicates that the identifiability of these models with the current data is much higher than random cases (25%). The recovery rate of the RSQL model was low (40%). Therefore, we concluded that appropriate QL, utility, and surprise models can be selected for each participant. However, as a cautionary note regarding the surprise model, the selection rate of the RSQL model for the simulated surprise model was slightly high (24%), and vice versa, which indicates that these models are difficult to distinguish from one another.

### Posterior predictive check

We conducted a multiple regression analysis of real preference data with simulated data from all models and found that the interaction of the group and quadratic term of age was significant in three models (utility: *β*=0.152, *p*=0.001, RSQL: *β*=0.122, *p*=0.020, surprise: *β*=0.135, *p*=0.021) (S10 Table). Therefore, we performed a correlation analysis between the simulated risk preference in these three models and each participant’s actual risk preference, and these models showed a similarly strong correlation with the actual choices (utility: *r*=0.83, RSQL: *r*=0.87, surprise: *r*=0.82) (S15 Fig.). These results indicate that these models can capture data sufficiently well to predict real behaviors.

Additionally, we performed the same analyses on the stay probability of sure choices, risky choices after a non-rewarding outcome, and risky choices after a rewarding outcome. For simulated sure choices, the interactions between the group and quadratic term of age were significant only in the utility (*β*=-0.055, *p*<0.001) and surprise (*β*=-0.071, *p*=0.020) models but not in the QL (*β*=0.001, *p*=0.938) and RSQL (*β*=-0.054, *p*=0.070) (S11 Table and S16a-d Fig.) models. In the utility and surprise models, the correlations between the simulated and real stay probabilities of sure choices were similarly strong (utility: *r*=0.70, surprise: *r*=0.68). However, the interaction for simulated risky choices after a non-rewarding outcome (S12 Table and S16e-h Fig.) and for simulated risky choices after a rewarding outcome (S13 Table and S16i-l Fig.) was not significant in any model.

### Supplemental analysis with open data

As a result of model comparisons using the data of Rosenbaum et al. (17) who found a nonlinear developmental change in risk preference in NTP participants of similar age as that in this study and used a similar task but with a different monetary context, we confirmed that the surprise model had a better fit based on the model evidence (S17a Fig.) and protected exceedance probability (S17b Fig.) than the other models. Accordingly, adding evidence to this study, the surprise model can explain the risk preference data well, including from the developmental perspective. Furthermore, with the regression analysis, we confirmed a significant nonlinear relationship between the quadratic term of age and surprise parameter (*β*=-0.228, *p*=0.006) (S18 Fig.).

## Discussion

This is the first study to investigate the nonlinear trajectory of age-related changes in risk preference in AUT and NTP participants and propose the underlying computational mechanism that best explains the risk preference. Contrary to our hypothesis, we did not find a group difference in the mean risk preference. Instead, we found a significant difference in the relationship between risk preference and the quadratic term of age between the AUT and NTP groups. Risk preference in the NTP group increased toward adolescence and decreased later, in line with the results of previous studies investigating different contexts (1–5). By contrast, the AUT group showed a developmental curve in the opposite direction: risk preference decreased toward adolescence and increased afterwards. Critically, the estimated parameter from the surprise model, the better-fitting model based on model comparison, revealed that the preference for surprise underlies the opposite patterns of developmental change in risk preference between the AUT and NTP groups.

Adolescence is a time of dramatic emotional and social change (33, 34), which also poses vulnerabilities to physiological and neural development (35, 36) in both NTP and AUT individuals. Although no studies have explicitly investigated the nonlinear trajectory of age-related change in risk preference in AUT, South et al. (37) reported the risk preference of AUT and NTP children and adolescents, which partly aligns with the current study. Their findings implied that risk preferences were similar between NTP and AUT adolescents, but there was a higher risk preference in AUT individuals than in NTP individuals during childhood. This result indicates that age is an important moderator of risk preference differences between AUT and NTP individuals.

For further interpretation of risk preference, the stay probabilities for each option helped us better evaluate the factors influencing risk preference in AUT. In the AUT group, regardless of whether the reward was acquired, the stay probability of risky choices decreased toward adolescence and increased afterwards, but it was the opposite for sure choices. These results indicate that among AUT youths, individuals who subjectively prefer risk choose the risky option regardless of the preceding objective outcome value, which may originate from the conformity of both positive and negative prediction errors, as a surprise. Additionally, this finding is consistent with AUT features and preferences for repetitive behavioral patterns. In the NTP group, individuals who subjectively avoided risk tended to stay with the sure choice, but with increasing age, the objective outcome value affected their stay probability for risky choices. Accordingly, in both groups, it seemed that risk-related choices were weighted not only by the objective outcome value but also by subjective value processing.

To uncover the computational mechanisms contributing to these behavioral indicators, we conducted computational modeling using reinforcement learning models that incorporated possible additional factors that could account for risk preference. The model comparison showed that the best-fitting model was the surprise model, which incorporated the surprise parameter that alters the reward sensitivity such that a larger prediction error further attenuates the reward value. Moreover, the relationship between the surprise parameter and quadratic term of age showed a significant group difference in the opposite pattern to that of the relationship between risk preference and the quadratic term of age. Furthermore, these results were supported by those of an additional analysis of the data of a previous risk-taking study on NTP youth (17) that found that the surprise model had a better fit than previous winning models, such as the RSQL and utility models. We also found a significant nonlinear relationship between the quadratic term of age and surprise parameter. These findings indicate that the preference for surprise is one of the key computational mechanisms underlying developmental changes in risk preference. Adolescence is often considered a developmental period with heightened sensitivity to rewards, resulting in risky behaviors (1–5). Such reward sensitivity may be represented by the surprise parameter, which reflects individual differences in confidence regarding surprise in decision outcomes.

Finally, these two findings were inconsistent with our initial hypothesis. First, AUT participants were not risk averse, as in previous studies with financial risk preference tasks targeting AUT adults (13–15), indicating that the target age and context of the task (i.e., presence or absence of financial reward) may be important factors for risk preference. Another finding that did not support our hypothesis was the preference for surprise. In our previous study (18), we assumed that surprises negatively affected the outcome value. However, in the present study, which used a game-like risk preference task, approximately half of the participants in both groups preferred the risky option, and the estimated surprise parameter was below zero, suggesting that surprises increased the value of the outcome for these participants. It is known that some individuals who self-report that they like surprises prefer mysterious consumption——the opportunity to be surprised——over non-mysterious consumption of equal expected value (38). Accordingly, risk preference may differ in different contexts and individuals from the perspective of surprise preference. Future studies should investigate developmental changes in risk preference in different contexts, such as financial or game-like tasks.

### Limitations

Our study is not without limitations. First, our target sample size was over 60 participants in each group similar to that of Rosenbaum et al. (17). However, after multiple data exclusion steps, data from only 60% of participants could be included in the final sample. Using these criteria, we confirmed that the accuracy increased as the task proceeded, similar to Rosenbaum et al. (17). Therefore, our results from the remaining data are rigid, but it is desirable to conduct an experiment without data loss. Another limitation was the variability of participant characteristics between groups, such as age and IQ, which were significantly different between the groups. Although these parameters, age and accuracy instead of IQ, were controlled for as covariates in the multiple linear regressions, future studies in a more controlled participant group are required.

### Conclusion

In conclusion, the current study is the first to demonstrate a significant difference in nonlinear developmental changes in risk preference between AUT and NTP participants. In adolescence, risk preference was similar between the AUT and NTP groups, but the opposite was true in childhood and adulthood. Using a computational modeling approach, we revealed the underlying mechanism of risk preference from the perspective of surprise preference. These findings indicate that in NTP individuals, adolescence is a developmental period in which risk is preferred because of the lowest aversion to surprise, whereas in AUT individuals, adolescence is a developmental period in which risk is avoided because of the highest aversion to surprise.

## Materials and Methods

### Participants

This study was conducted as part of a multiple-study protocol project targeting students or alumni of a private school, Musashino Higashi Gakuen, including both AUT and NTP participants. Individuals had been clinically diagnosed with autism spectrum disorder by at least one pediatrician, child psychiatrist, or clinical psychologist. In this study, 125 participants aged 6–30 years completed an online task in their homes. After multiple data exclusion steps, we analyzed the data of 75 participants (17 females, 57 males, and 1 person who did not prefer to report; mean age = 16.03 years, standard deviation = 5.86; 31 in the AUT group and 44 in the NTP group) (Table S1). The parents of all participants completed the Japanese version of the Social Communication Questionnaire (SCQ; Rutter et al., 2003). Among these participants, 27 and 25 in the AUT and NTP groups, respectively, reported their intelligence quotient (IQ) scores measured for previous projects (e.g. Asada et al., 2024; Kikuchi et al., 2022; Akechi et al., 2018) using either the WISC-III or WAIS-R.

All participants and their parents were given instructions, and they completed an informed consent form before study participation. This study adhered to the Declaration of Helsinki and was approved by the Committee on Ethics of Experimental Research on Human Subjects, Graduate School of Arts and Sciences, University of Tokyo (156-17).

### Exclusion criteria

We first excluded data from eight participants who did not press the button in >10% of the trials. We then screened the data for a reaction time that was too fast (mean of all trials <300 ms) and the same button being pressed in >90% of the trials; these applied to none of the participants. Additionally, based on the study by Rosenbaum et al. (17), we excluded the data of 42 participants whose mean accuracy in the test trials in the second and third blocks was <60%. After applying these criteria, we used regression analysis to confirm that participants learned the probability of each option block-by-block (t=6.065, p<0.001) (mean accuracy of test blocks: 1=0.63, 2=0.77, 3=0.79).

### Experimental design

PsychoPy v2021.2.3, presented using Pavlovia (pavlovia.org), was used to build the online task. The majority of participants completed the task on their personal computers (PC), and the remainder completed the task on their tablets (PC: 62 participants, tablets: 13 participants) (S1 Table). The stimuli used in the task (aliens, treasures, and backgrounds) were adapted from open-access materials created by Kool et al. (39), whereas the other stimuli (rockets) were adapted from an online site (iStockphoto.com).

### Treasure task

In this study, we created a treasure task based on the risk-sensitive reinforcement learning task of Rosenbaum et al. (17) (Fig. 1). As a cover story, the participants rode a rocket to a planet and received treasures from an alien who gave them 0–8 treasures. The number of treasures the participants received depended on the rocket they chose, and their challenge was to collect as many treasures as possible and obtain the large rocket. There were five rockets including three deterministic rockets for 0, 2, and 4 treasures, respectively, and two probabilistic rockets for 0 or 4 and 0 or 8 treasures, respectively (50% each). In the task, there were three blocks and 183 trials, including 42 sure vs. risky choices (24 choices for a 100% 2-treasure rocket vs. a 50% 0- or 4-treasure rocket and 18 choices for a 100% 4-treasure rocket vs. a 50% 0- or 8-treasure rocket), 24 other choices (a 100% 2-treasure rocket vs. a 50% 0- or 8- treasure rocket), 42 test choices that ensured that participants learned the features of each rocket (e.g., a 100% 2-treasure rocket vs. a 100% 4-treasure rocket), and 75 forced-choice trials in which participants were forced to learn the features of each rocket. The trial order was pseudo-randomized based on seven templates created from the order of the seven participants in the study by Rosenbaum et al. (17).

### Basic analysis

#### Risk preference

First, we compared group differences in risk preference using a t-test. Subsequently, using multiple regression analysis, we examined whether there was a group difference in risk preference that changed nonlinearly with age by modeling the interaction term of the quadratic term of age and groups. As confounding covariates, in addition to the reported gender (male, female, or preferred not to report) and execution device (PC or tablet), we modeled the accuracy of the test trials in the second and third blocks, which was significantly correlated with IQ scores among participants who reported their IQ score (r=0.34, p=0.012), to remove the effect of the intellectual component associated with this task. Models were assessed using the lm_robust command from the “estimatr” package (40) with heteroskedasticity-consistent 0 robust standard errors. This method estimates the standard errors under heterogeneous variances and applies a weighting function to reduce the influence of outlier data. In these analyses, categorical variables were coded as 1 = AUT and −1 = NTP for the group and as 1 = PC and −1 = tablet for the device; continuous variables—age and accuracy—were scaled to mitigate multicollinearity issues. We checked for the multicollinearity of regressors within a model using the “performance” package (41).

Furthermore, as a consecutive analysis to confirm the nonlinear developmental change in risk preference in each group, we conducted a multiple regression analysis for risk preference within each group by using the model with the quadratic term of age, in addition to linear term of age, reported gender, execution device, accuracy of the test trials, and SCQ score.

#### Stay probability

We calculated the stay probability (i.e., the probability of choosing the same option consecutively) of the sure and risky choices both after the rewarding and non-rewarding outcomes. In these calculations, we did not consider the outcome of the forced-choice trials that appeared between the re-risk choice trials. We investigated the group differences in each stay probability that changed nonlinearly with age by modeling the interaction term of the quadratic term of age and groups. The other regressors were the same as those used in the analysis for risk preference. Furthermore, as a consecutive analysis to confirm the nonlinear developmental change in each stay probability in each group, we conducted a multiple regression analysis within each group using the same regressors as in the analysis for risk preference.

### Computational modeling

#### Model description

In this study, we fit three widely used models—the QL, utility, and RSQL models—and our proposed model, the surprise model. For each model, the learning rate was constrained to the range, 0 ≤ *α*, *α*^+^, *α*^−^ ≤ 1, with a beta (2,2) prior distribution, and the inverse temperature was constrained to the range, 0 ≤ *β* ≤ 20, with a gamma (2,3) prior distribution. The utility parameter was constrained to the range, 0 ≤ *ρ*≤ 2.5, with a gamma (1.5,1.5) prior distribution. Additionally, the modulation rate of the surprise model was constrained to −1 ≤ *d* ≤ 1 with a uniform prior distribution.

#### QL model

The QL model is used as the base model for the other three models. The QL model incorporates the Rescorla-Wagner rule, where only the Q-value of the chosen option is updated based on a prediction error that explains the observed behavior by computing the action value Q(*t*) for each trial *t*, which represents the expected outcome of the action. The Q-value of the chosen action is iteratively updated based on a prediction error, which is the difference between the expected outcome Q(*t*) and received outcome *r*, by a learning rate *α*.

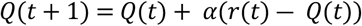

#### Utility model

The utility model is a QL model that incorporates nonlinear subjective utilities for different amounts of rewards. In this model, the reward outcome is exponentially transformed by *ρ*, which represents the curvature of the subjective utility function for each individual.

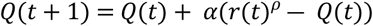

#### RSQL model

In the RSQL model, which is a QL model, positive and negative prediction errors have asymmetric effects on learning. Specifically, there are separate learning rates: *α*^+^ and *α*^−^ for positive and negative prediction errors, respectively.

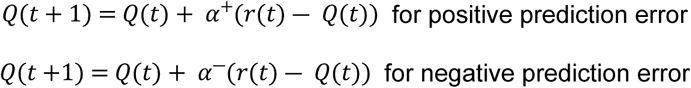

#### Surprise model

In the surprise model, the received outcome *r* is affected by surprise (absolute value of the prediction error). In this model, *S*(*t*) is the subjective utility modulated by surprise. The degree of modulation is controlled by the parameter *d* as follows:

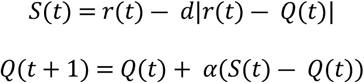

For all models, the probability of choosing option *i* during trial *t* is provided by the softmax function:

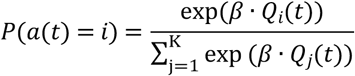

where *β* is the inverse temperature parameter that determines the sensitivity of the choice probabilities to differences in the values, and *K* represents the number of possible actions (in the present study, *K* = 2); moreover, a(*t*) denotes the option chosen in trial *t*.

### Parameter estimation

We fit the parameters of each model using the maximum a posteriori (MAP) estimation, which improves parameter estimates by incorporating prior information on parameter values (27, 42). We also approximated the log marginal likelihood (model evidence) for each model using the Laplace approximation (43). We used the R function “solnp” in the “Rsolnp” package (44) to estimate the fitting parameters.

### Model comparison

For model selection, the model evidence (log marginal likelihood) for each model and participant was subjected to random-effects Bayesian model selection (BMS) using the “spm_BMS” function in SPM12 (45). BMS provides a less biased and statistically more accurate way to identify the best model at the group level by estimating the protected excess probability, which is defined as the probability of a particular model being more frequent in the population among a set of candidate models (27). We conducted a model comparison for all participants in each group. To visualize how well the winning model fit the data, we also determined the number of participants that best fit each model (27).

### Estimated parameters

After the model comparison, we investigated group differences in the estimated parameters of the winning model, which changed nonlinearly with age, by modeling the interaction term of the quadratic term of age and groups. The other regressors were the same as those used in the risk preference analysis.

Furthermore, as a consecutive analysis to confirm the nonlinear developmental change in the estimated parameters in each group, we conducted a multiple regression analysis within each group with the same regressors as in the risk preference analysis.

### Parameter recovery

We validated the fitting procedure for the task using parameter recovery. We conducted parameter recovery to assess the reliability of parameter estimation procedures; specifically, we assessed how accurately parameters were estimated when the true generative model and its parameter values were known (27, 46, 47). For each model, we simulated the choice data for the same experimental paradigm as the treasure task using one of the seven templates for 1,000 agents, with the values of each parameter drawn randomly and uniformly from the range of possible parameter values as real data. Only for the inverse temperature, to avoid convergence errors, we constrained the data in all models to the range, 0.05 ≤ *β* ≤ 10. We then fit the simulated data with the MAP estimation. We tested the recoverability of the model parameters by checking the correlation between the parameters that generated the data and those generated by the model fit.

### Model recovery

To test the discriminability of each model using the simulated data for parameter recovery, we fitted all four models with MAP estimation to all the simulated data from these models, including those not used to generate the data for parameter recovery. Model recovery refers to the ability to correctly identify a true data-generating model using model comparison techniques (27, 46, 47). We then used log marginal likelihood to assess whether the data-generating model was also the best-fitting model for each participant.

### Posterior predictive check

We performed a posterior predictive check that analyzed the simulated data in the same way as the analyses of empirical data to validate that each model adequately captured behavioral data (27, 28). Using each participant’s estimated parameters, we simulated choices with the same task, one of seven templates, for 10,000 agents with each model and recorded the median proportion of risk preference across 10,000 agents. Using the simulated risk preference data from all four models, we conducted a multiple regression analysis with the same regressors as those in the real behavioral data. We then conducted a correlation analysis between simulated risk preference and real risk preference only in the models where the interaction of group and the quadratic term of age was significant for simulated risk preference. For additional analyses, we calculated the stay probability from the simulated data and performed the same analysis as that for risk preference.

### Supplemental analysis with open data

As the task used in this study was based on the study by Rosenbaum et al. (17), to confirm that the surprise model was better fitted to the risk preference data with similar age diversity, we conducted model fitting and model comparisons. For model fitting, the parameters were set to be the same as those used in this study. Additionally, to confirm the nonlinear relationship between age and the surprise parameter, we conducted a regression analysis of the surprise parameter using the quadratic term of the scaled age.

## Acknowledgments

We would like to acknowledge all the participants, their families, and the teachers at Musashino Higashi Gakuen.

## Open Practices Statement

The experimental task and all codes used in the analysis are available from the Open Science Framework (https://osf.io/b5rt2/). Data are available upon request owing to privacy/ethical restrictions.

## Abbreviations

AUT: autistic
BMS: Bayesian model selection
MAP: maximum a posteriori
NTP: neurotypical
RSQL: risk-sensitive Q-learning
QL: Q-learning
VIF: variance inflation factor

## Supporting Information captions

**S1 Text. Supplemental Discussion**: **Learning rate and inverse temperature.**

**S2 Text. Model comparison.**

**S1 Table. Demographic data.** Note: IQ* for only participants who reported the scores. Gender, M = male, F = female, N = not preferred; IQR, interquartile range; SCQ, Social Communication Questionnaire; AUT, autism; NTP, neurotypical

**S2 Table. Regression table for risk preference and the surprise parameter.** The value inside of the () is the p value. Group (AUT = 1, NTP = −1), device (PC = 1, Tablet = −1), gender (M = male, F = female, N = not preferred to report). AUT, autism; NTP, neurotypical

**S3 Table. Regression table for risk preference in each group.** The value inside of the () is the p value. Group (AUT = 1, NTP = −1), device (PC = 1, Tablet = −1), gender (M = male, F = female, N = not preferred to report). AUT, autism; NTP, neurotypical; SCQ, Social Communication Questionnaire

**S4 Table. Regression table for the surprise parameter in each group.** The value inside of the () is the p value. Group (AUT = 1, NTP = −1), device (PC = 1, Tablet = −1), gender (M = male, F = female, N = not preferred to report). AUT, autism; NTP, neurotypical

**S5 Table. Regression table for parameters within the surprise model.** The value inside of the () is the p value. Group (AUT = 1, NTP = −1)

**S6 Table. Regression table for stay probability.** The value inside of the () is the p value. Group (AUT = 1, NTP = −1), device (PC = 1, tablet = −1), gender (M = male, F = female, N = not reported). AUT, autism; NTP, neurotypical

**S7 Table. Regression table for stay probability in each group.** The value inside of the () is the p value. Group (AUT = 1, NTP = −1), device (PC = 1, tablet = −1), gender (M = male, F = female, N = not reported). Sure, stay probability of a sure choice; stay_NR, stay probability of a risky choice after a non-rewarding outcome; stay_R, stay probability of a risky choice after a rewarding outcome; AUT, autism; NTP, neurotypical

**S8 Table. Best fitted model for each participant.** AUT, autism; NTP, neurotypical

**S9 Table. Recoverability of parameters in each model.**

**S10 Table. Posterior predictive check for risk preference.** The value inside of the () is the p value. Group (AUT = 1, NTP = −1), device (PC = 1, tablet = −1), gender (M = male, F = female, N = not reported). AUT, autism; NTP, neurotypical

**S11 Table. Posterior predictive check for sure choices.** The value inside of the () is the p value. Group (AUT = 1, NTP = −1), device (PC = 1, tablet = −1), gender (M = male, F = female, N = not reported). AUT, autism; NTP, neurotypical

**S12 Table. Posterior predictive check for risky choices after a non-rewarding outcome.** The value inside of the () is the p value. Group (AUT = 1, NTP = −1). AUT, autism; NTP, neurotypical

**S13 Table. Posterior predictive check for risky choices after a rewarding outcome.** The value inside of the () is the p value. Group (AUT = 1, NTP = −1), device (PC = 1, tablet = −1), gender (M = male, F = female, N = not reported). AUT, autism; NTP, neurotypical

**Fig. S1. Variance inflation factor in the multiple regression analysis of (a) risk preference and (b) surprise parameter.**

**Fig. S2. Model coefficients for risky choices in each group.** Parameters within models were coded as group (AUT = 1, NTP = −1), device (PC = 1, Tablet = −1), and gender (M = male, F = female, N = not reported). AUT, autism; NTP, neurotypical; SCQ, Social Communication Questionnaire

**Fig. S3. Variance inflation factor in the multiple regression analysis of risky choices in each group.**

(a) AUT and (b) NTP.

AUT, autism; NTP, neurotypical

**Fig. S4. Model coefficients for the surprise parameter in each group.** Parameters within models were coded as group (AUT = 1, NTP = −1), device (PC = 1, Tablet = −1), and gender (M = male, F = female, N = not reported).

AUT, autism; NTP, neurotypical; SCQ, Social Communication Questionnaire

**Fig. S5. Variance inflation factor of the surprise parameter in each group in the multiple regression analysis. (a)** AUT and **(b)** NTP.

AUT, autism; NTP, neurotypical

**Fig. S6. Evaluation of (a) relationship between age and estimated alpha and (b) relationship between age and estimated beta.** The regression line is derived from a linear regression model that includes both linear and quadratic age terms. The data points represent individual participants. The lines and 95% confidence intervals were estimated under heterogeneous variances to reduce the influence of outlier data.

AUT, autism; NTP, neurotypical

**Fig. S7. Evaluation of (a) alpha and (b) beta model coefficients.** Parameters within models were coded as group (AUT = 1, NTP = −1).

AUT, autism; NTP, neurotypical

**Fig. S8. Variance inflation factor in the multiple regression analysis of the estimated parameters (a) alpha and (b) beta.**

**Fig. S9. Variance inflation factor in the multiple regression analysis of the stay probability of (a) sure choices, (b) risky choices after a non-rewarding outcome, and (c) risky choices after a rewarding outcome.**

**Fig. S10. Model coefficients for each stay probability in each group.** Parameters within models were coded as device (PC = 1, Tablet = −1) and gender (M = male, F = female, N = not reported).

AUT, autism; NTP, neurotypical; SCQ, Social Communication Questionnaire

**Fig. S11. Variance inflation factor in the multiple regression analysis of the stay probability of sure choices in (a) the AUT and (b) NTP groups, risky choices after a non-rewarding outcome in the (c) AUT and (d) NTP groups, and risky choices after a rewarding outcome in the (e) AUT and (f) NTP groups.**

AUT, autism; NTP, neurotypical; SCQ, Social Communication Questionnaire

**Fig. S12. Model comparison based on evidence of each model (log marginal likelihood) in the (a) AUT and (c) NTP groups and Bayesian model comparison in the (b) AUT and (d) NTP groups.**

AUT, autism; NTP, neurotypical

**Fig. S13. Distribution of estimated parameters in the surprise model.**

ASD, autism spectrum disorder; NTP, neurotypical

**Fig. S14. Results of model recovery.** The confusion matrix represents the frequency with which data generated with each model were best fit by each model.

**Fig. S15. Correlation between real risk preference and simulated data in each model.**

**Fig. S16.** Correlation between the stay probability and simulated data in each model. The (a-d) sure choices, (e-h) risky choices after a non-rewarding outcome, and (i-l) risky choices after a rewarding outcome were evaluated.

**Fig. S17. Model comparison using data from Rosenbaum et al.** (**2022**) **based on (a) evidence of each model (log marginal likelihood) and (b) Bayesian model comparison.**

**Fig. S18. Relationship between risk preference and age based on data from Rosenbaum et al.** (**2022**). The regression line is derived from a linear regression model that includes both linear and quadratic age terms. The data points represent individual participants. The lines and 95% confidence intervals were estimated under heterogeneous variances to reduce the influence of outlier data.

